# Stable and Reversible Functionalization of the Native Phosphate Groups on Live Cells

**DOI:** 10.1101/462044

**Authors:** Joydeb Majumder, Gaurav Chopra

## Abstract

Surface modification of live cells has important biomedical and therapeutic applications. Current methods to label cells require artificial cell surface engineering (via metabolic, docking or anchoring methods) before conjugative chemistries, which is not always trivial to accomplish and/or not appropriate for multiple cell types. A new method without the need of initial cell surface anchoring will greatly facilitate live cell surface labelling. Herein, we provide a general strategy for live cell functionalization that utilizes the native phosphate groups on every cell. We have designed a dual conjugation cargo molecule with a cationic side chain for non-covalent bonds with the negatively-charged cell surface and a phosphoric acid containing ligand for covalent bonding with the cell membrane phospholipid phosphate. Our dual conjugation strategy on live cell surfaces is non-toxic with enhanced stability to functionalize live cells. This provides a stable, reversible and reusable reagent with direct conjugation strategy to image live cell membranes.

**Significance:** The ability to label live cell surfaces has many applications ranging from *in vivo* monitoring of cell populations to diagnostics and use of cells as drugs. Thus far, most reported strategies to label cell surfaces are not broadly applicable or easy to use for any cell type as it has relied on engineering cells with artificial moieties or conjugations that may affect cellular function. We provide a general solution to this long-standing problem by developing two-sided functionalization of the phosphate moieties that are ubiquitous on all cells. We show one application of our chemical strategy as a general-purpose live-cell membrane imaging reagent with long-time stability. Our strategy is broadly applicable to imaging, sensing, drug delivery, bioengineering, diagnostics and cell therapy.

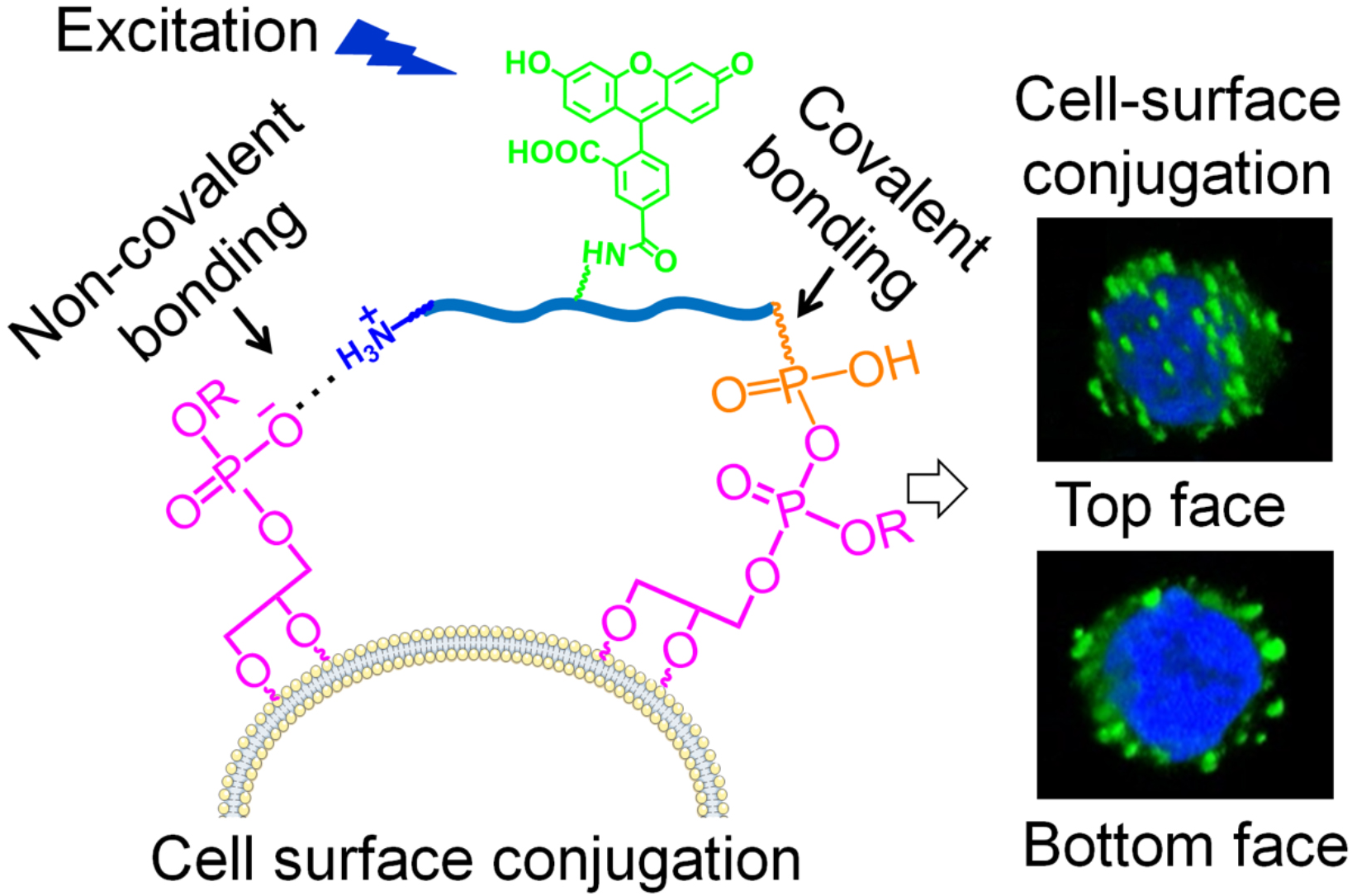

## Introduction

Surface modification of live cells has many biological applications including imaging, control of cell surface interactions, tracking and sensing biological environments in vitro and in vivo.^1,2^ Recently, cell surface functionalization has received significant attention from researchers and clinicians perhaps due to several biomedical applications.^3,4^ Several methods have been developed to functionalize the cell surface with cargo vehicles and therapeutic agents. Most popular conjugation methods include hydrophobic anchoring,^5–13^ chemo-selective conjugation,^14–18^ PEGylation^19^ etc. These methods are generally based on the conjugation of various functional groups with the cell surface ligands or proteins via (1) electrostatic interactions between positively-charged polyelectrolytes and negatively-charged cell surfaces such that negatively charged cell membrane bind with positively charged polycationic molecules including poly-peptides with basic amino acid side chain or cationic polyelectrolytes;^20^ (2) hydrophobic interactions between polyelectrolyte backbones and cell membrane’s lipid bilayers;^21^ and (3) covalent interactions with proteins and functional groups on cell surfaces, for example, cargo molecule containing N-hydroxysuccinimide ester (NHS) groups are widely used to form covalent bonds with the amino groups of membrane proteins.^22,23^ Chemical ligations on cell surfaces have been also achieved by using unnatural functional groups.^24^ Recently, there have been major advances to modify live cell surface with synthetic functional polymers,^25^ macromolecular crowding,^26^ and conjugation of well-engineered therapeutic loaded nanoparticles^27–31^ using artificial anchors on the cell membrane for cell therapy applications.

However, live cell surface modification done till date show limited time stability and may affect their desired function. For example, the overall stability for PEG-lipids conjugation on cell surface was no longer than 1-2 days before all the PEG molecules dissociated from the cell surface.^32^

Furthermore, incorporating hydrophobic chain in the polymer scaffold results in uptake by the cells affecting normal cell function. On the other hand, metal-free covalent conjugation of live cells by chemo-selective reactions on transmembrane proteins provide slightly longer stability of the surface conjugation^33–35^ but results in toxic effects and disruption of normal cellular function^36^ due to non-specific binding of the cargo molecules with other amine groups on membrane proteins. Finally, there are also limitations in the development of nanoparticles-based cell-conjugates for in vivo cell therapy. For example, nanoparticles cause toxic side effects after entrapment in the reticuloendothelial system of the liver and spleen,^37^ and bio-compatibility of synthetic nanoparticles made from inorganic materials raises a major concern for translation.^38^

Live cell chemical conjugation is a research area where major challenges and problems^39^ remain to widely adopt established methods in practice. Development of new strategies for stable conjugation of therapeutics/cargo for live cell surface will result in better viability and function of these modified cells. Keeping both the stability and viability objectives in mind, here we provide a general cell surface conjugation strategy using the native phosphate groups on cell membrane. We have decorated one side of the molecule with a cationic functionality which will form noncovalent interaction with the negatively charged cell surface and the other side of the cargo molecule with a phosphoric acid containing ligand such as adenosine di-phosphate (ADP) which will facilitate phospho-ester covalent bonds under physiological condition with the cell surface phospholipid phosphate functionality. Our dual chemical conjugation approach provides longer stability of the cell-surface conjugation and shows no effect on viability due to the formation of the natural phospho-ester bond instead of unnatural chemical reactions.^24^

## Results and Discussion

### Cargo design, synthesis, and phospholipid bond formation

First, we wanted to see the effect of small-molecule fluorophore conjugates on cells. We synthesized an ADP-fluorescein conjugate and treated Jurkat T cells. Similar to fluorescein treatment, ADP-fluorescein conjugate was internalized by T cells (Supplementary Figure S1). This led to the design of a cargo molecule similar to a polymer/macromolecule carrier. We designed and synthesized a cargo backbone with only side chain cationic group (non-covalent cargo) while another cargo backbone with both side chain cationic and phosphate groups (covalent cargo, Figure 1a-b, Supplementary Figure S2). We hypothesized that phosphate anions of phospholipids on cell membranes form phospho-ester bonds with phosphoric acid group on ADP via condensation reaction. To test our hypothesis, we also synthesized phospholipid linked non-covalent and covalent cargos (Figure 1c, Supplementary Figure S3) and characterized by NMR and FT-IR spectroscopy (Supplementary Figure S15-23). For phospholipid fatty acid, P=O stretching band appeared at 1170 cm^-1^, and P=O stretch was at 1156 cm^-1^ for noncovalent cargo (Supplementary Figure S22). The decrease in P=O stretching band indicated its presence as anion and/or conjugated form such as phospho-ammonium group. Similarly, for covalent phospholipid cargo, a larger decrease in P=O stretching band suggests a covalent bond in extended conjugated form as a tri-phosphoester bond. The decrease in P=O stretching band was observed at 972 cm^-1^ for covalent phospholipid, compared to 1103 cm^-1^ in ADP and 1075 cm^-1^ in the covalent cargo (Supplementary Figure S23).

**Figure 1.**
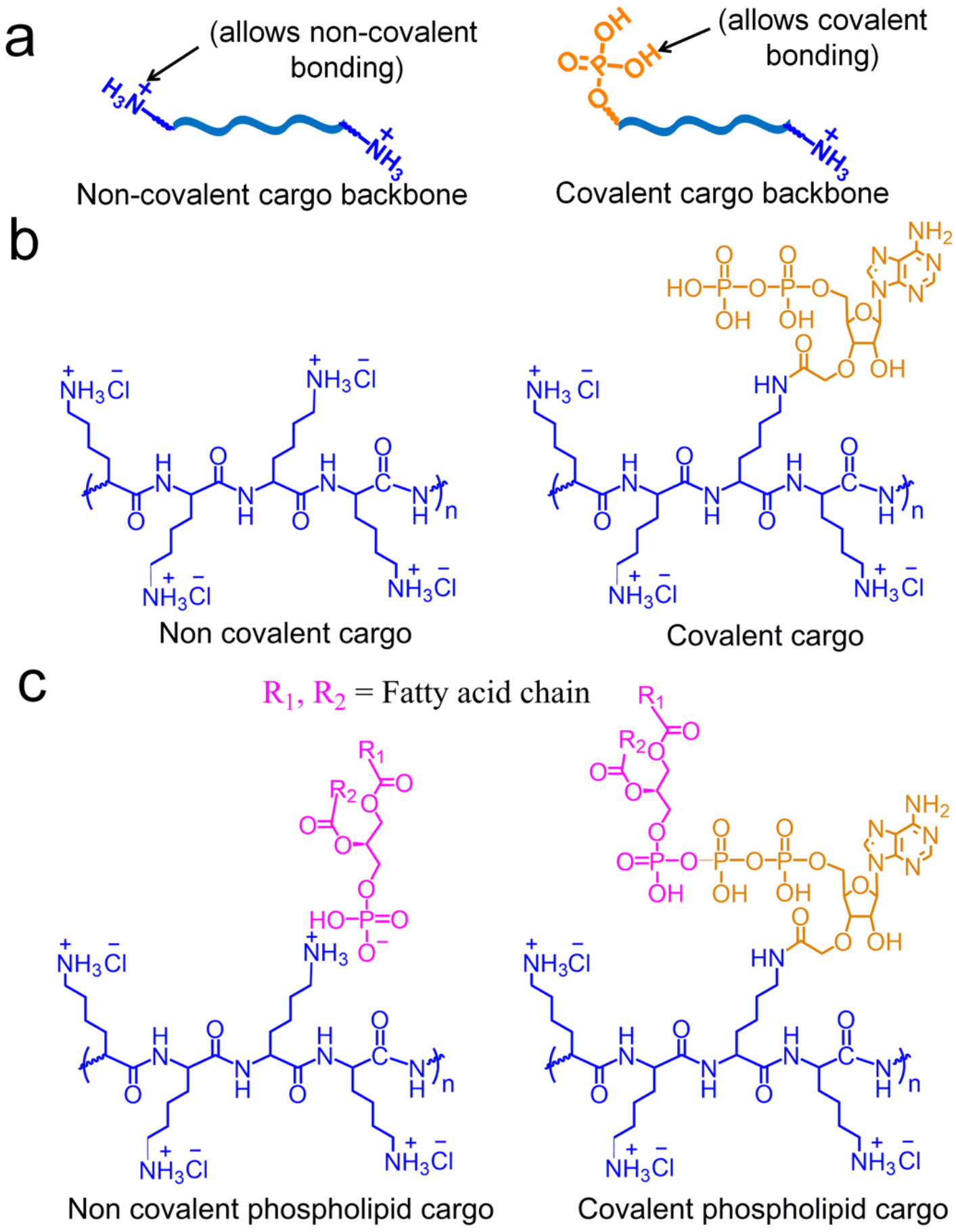
Design strategies and structures of cargo molecules for live cell surface chemical conjugation. (a) Design of the noncovalent and dual-conjugated covalent cargo backbone with both cationic side chain and phosphoric acid functionality. (b) Structures of the non-covalent and covalent cargo molecule, and (c) phospholipid linked non-covalent and covalent cargo.

### Dual chemical conjugation of live cell-surface avoids cellular internalization

After establishing bond formation between our synthetic cargo molecules and the phospholipid, we hypothesized that our noncovalent and dual covalent cargo molecules will interact with the native cell-surface phosphates, as outlined in Figure 2a. We conjugated a fluorophore molecule (fluorescein) with both non-covalent and dual conjugation covalent cargo molecules to locate the presence of the cargo molecules on cells (Figure 2b, Supplementary Figure S4). Next, we were interested to see whether the two cargo molecules will conjugate on the membrane of Jurkat T cells. We identified optimal conditions for the cargo (150 kD, Figure S5) and used it with an optimal concentration of 0.1 mg/mL to treat the cells at RT for 30 minutes on an orbital shaker (Supplementary Figure S6). Next, cells were stained with Hoechst 33342 and fluorescence confocal microscopy images of the Jurkat T cells are shown in Figures 2c and Supplementary Figure S7. Fluorescein is internalized by the cells as observed by perfect overlap of blue fluorescence of Hoechst 33342 and green fluorescence of fluorescein in the nucleus of the Jurkat cells (Figure 2c). When we used the fluorescein conjugated cargo, the blue fluorescence of Hoechst was observed in the nucleus and green fluorescence of fluorescein was observed at the periphery of the cells. The live cell surface conjugation efficiency of the of the dual conjugation covalent cargo was better than the non-covalent cargo. We did not observe internalization of non-covalent cargo or dual-conjugated cargo molecules but the latter molecule results in a brighter florescent signal suggesting that our hypothesis to develop a dual-conjugated cargo was correct to avoid cell surface internalization. Furthermore, to verify the need for dual cell conjugation, we synthesized a cargo by protecting its cationic ammonium side chains with BOC-groups. We observed no surface conjugation of the Jurkat T cells with the BOC-cargo clearly indicating the plausible role of the cargo decoration chemistry for cell-surface bond formations (Supplementary Figure S8).

**Figure 2.**
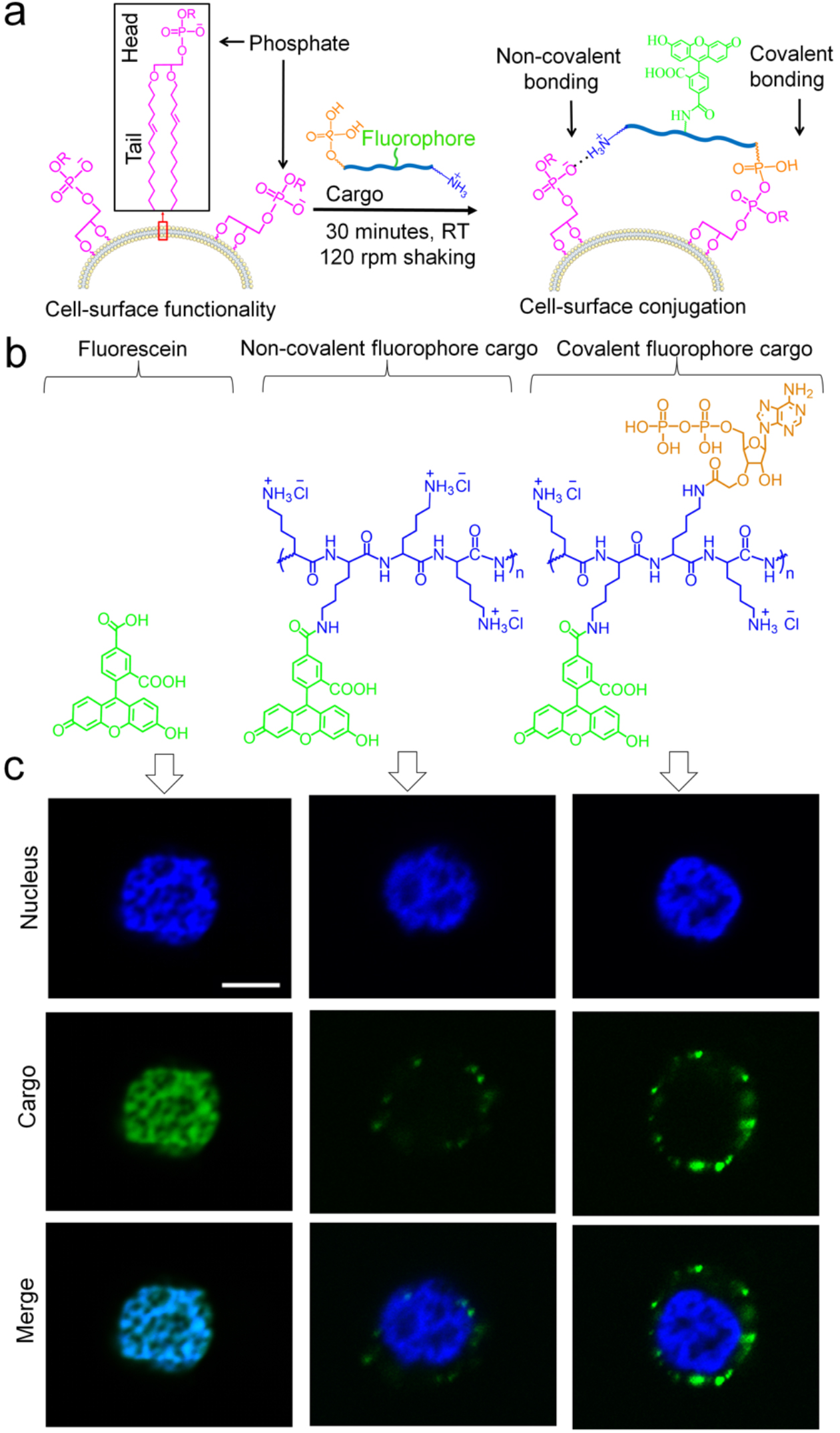
Chemical structures for live cell surface conjugation. (a) Cell surface functionality with negatively charged phosphate. Covalent and non-covalent conjugation of the fluorophore linked cargos on the cell surface. (b) Structures of the fluorescein, noncovalent and covalent cargos tagged with fluorescein. (c) Confocal images of the Jurkat T cells after treatment with fluorescein and fluorescein-tagged cargo molecules and followed by Hoechst 33342 nuclear staining. Both blue fluorescence of Hoechst 33342 and green fluorescence of fluorescein were observed in the nucleus of the Jurkat cells for the treatment of fluorescein molecule whereas green fluorescence of fluorescein was observed at the periphery of the Jurkat T cells for both the dual-conjugate covalent and non-covalent cargo treatments, indicating cell surface conjugation. Scale bar is 5 μm.

### Higher stability and low cytocompatibility of dual cell-surface conjugation on live cells

A major objective of this work is to develop cell-surface conjugation method with long time stability under physiological conditions (37 °C and pH 7.4) for several days. To our knowledge, the current stability of live cell conjugation, using artificial techniques that are hard to use for multiple cell types, is less than 72 hours.^32^ We incubated surface modified Jurkat T cells at 37 °C and pH 7.4 for the period of 6 days by simply adding our cargo molecules to the culture. The images were recorded after 1, 3 and 6 days (Figure 3a). Both the non-covalent and dual fluorescein tagged cargo molecules show stability to the surface of the Jurkat cells – as revealed by green fluorescence on the cell membrane after one day of conjugation but the dual covalent cargo molecule shows stability with more than 50% florescent remaining after six days in culture (Figure 3b). It is well known that activated T-cells results in lower surface redox levels with reduced number of membrane thiol (-SH) groups.^40,41^ We verified stability of our cargo with interleukin-2 (IL-2) activated Jurkat T cells (Supplementary Figure S9) suggesting possible use of conjugated immune cell therapy as future applications. This data clearly suggests a long-time stability of the surface conjugation for the dual covalent and non-covalent conjugation compared to only non-covalent cell conjugation.

**Figure 3.**
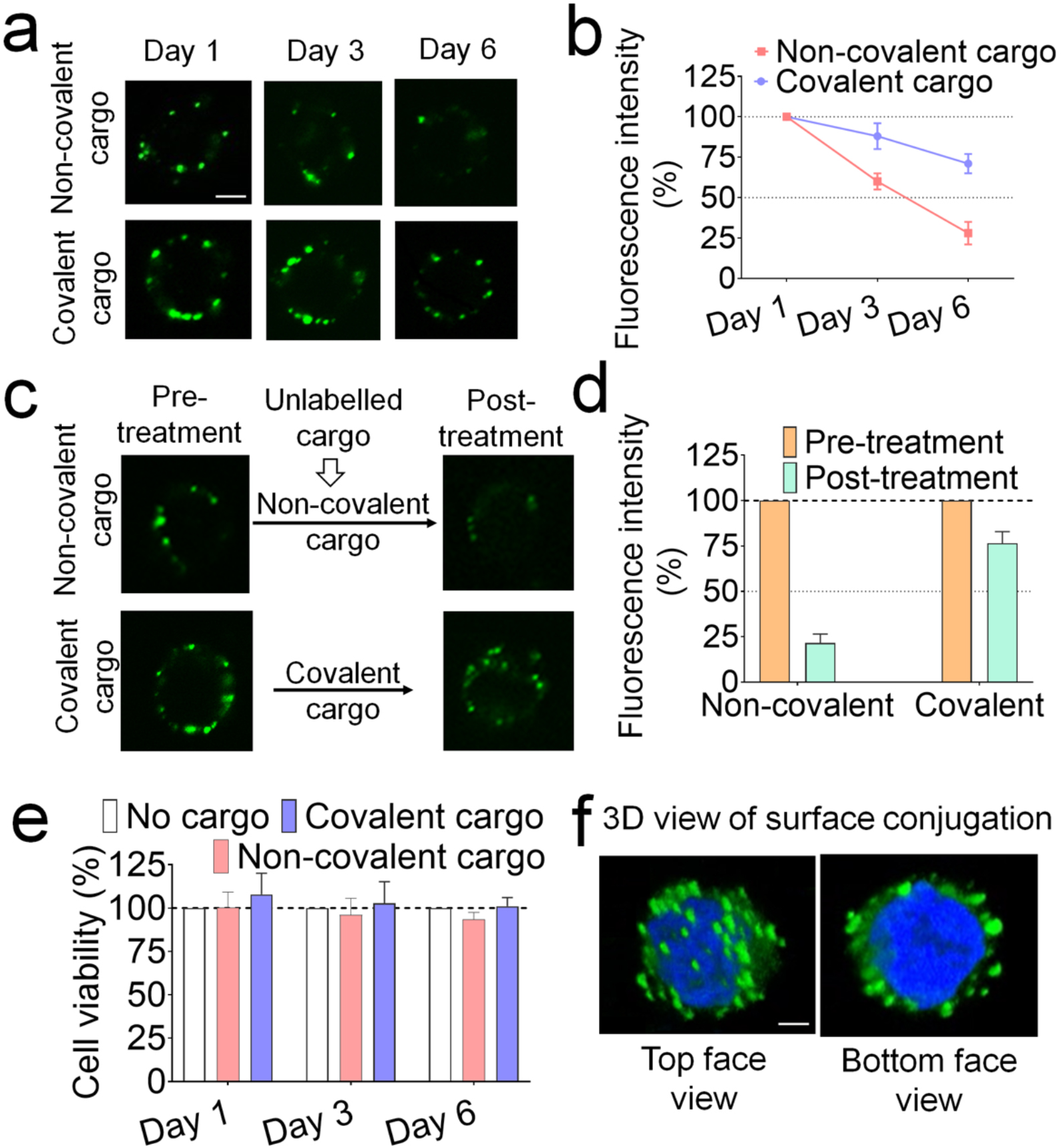
Stability and viability and of the surface modified Jurkat T cells. (a) Confocal laser microscopic images of the surface modified Jurkat T cells conjugated with both the cargos after 1, 3 and 6 days. (b) Quantification of surface fluorescence intensity of cells in Figure 3a. (c) Stability of the surface conjugated cells in presence of no fluorophore tag respective cargos. (d) Quantification of surface fluorescence intensity of cells in Figure 3c. (e) Viability of Jurkat T cells after their surface conjugation with the cargos. (f) 3D view of surface conjugated Jurkat T cells with covalent cargo. Dual-conjugation of the covalent cargo with Jurkat T cell is cytocompatible and stable under physiological conditions (37 ^°^C and pH of 7.4) over 6 days. Scale bar is 5 μm.

In order to investigate whether the phosphate bearing groups of the dual-conjugated covalent cargo have stable bonding with the cell surface functionality in a competing environment, we conjugated Jurkat T cells with fluorophore tag covalent cargo for 30 minutes and then washed with PBS. Next, the cells were treated with cargo molecules with no fluorophore tag (competing for similar bonding with florescent cargo molecules) for 30 minutes in growth media at 120 rpm at room temperature. We also performed the same experiment with non-covalent florescent cargo molecule in a competing presence of no fluorophore tagged non-covalent cargo (150 kD D-lysine chain polymer) for 30 minutes. The results indicate that the green fluorescence intensity on the surface of live cells conjugated with fluorophore-tag non-covalent cargo reduced significantly, compared to fluorophore-tag covalent cargo cell surface conjugation (Figures 3c-d, Supplementary Figure S10). Our data clearly indicates additional bonding interactions of the dual-conjugated covalent cargo, compared to the non-covalent cargo, with cell surface based on our cargo design strategies.

Keeping cells alive after cell surface conjugation is a requirement for future applications. To see the cytotoxic effect of surface conjugation, we performed viability test of the surface modified Jurkat T cells for 6 days. We performed the cell-titer blue viability assay resulting in more than 95% surface modified Jurkat T cells viability after 6 days (Figure 3e). Confocal 3D images of surface conjugated Jurkat T cell with the stable and non-toxic covalent fluorophore cargo is shown in Figure 3f.

### Cell-surface chemical conjugation for live cell membrane imaging

Based on the stability and non-internalization of the conjugated cargo on cell membranes, we were interested to validate the use of our florescent labelled covalent cargo as a reagent for membrane imaging of live cells. We treated multiple live cells to show the universal nature and usability of our live cell conjugation chemistry for membrane imaging applications. We treated, Jurkat T cells, human natural killer (NK-92) cells, mouse macrophage (RAW264.7) cells, human microglia (HMC-3) cells, and human prostate cancers (LNCaP, C4-2) cells with the dual-conjugated covalent cargo reagent for 30 minutes and observed membrane imaging of these live cells (Figure 4, Supplementary Figure S11). This is a useful application of our method since there are very limited and very expensive non-toxic staining reagent for live cell membrane imaging that do not show long-term stability. Our data clearly indicates that our dual cargo molecule is a suitable non-toxic reagent for any live cell membrane imaging applications. To further validate the usability of our reagent for membrane imaging of fixed cell we also treated our dual covalent cargo with fixed Jurkat T cells and observed enhanced membrane imaging of cells (Supplementary Figure S12) that is comparable to other fixed cells reagents. These results suggest that our stable and non-toxic conjugate reagent can be used to image both fixed and live cells.

**Figure 4.**
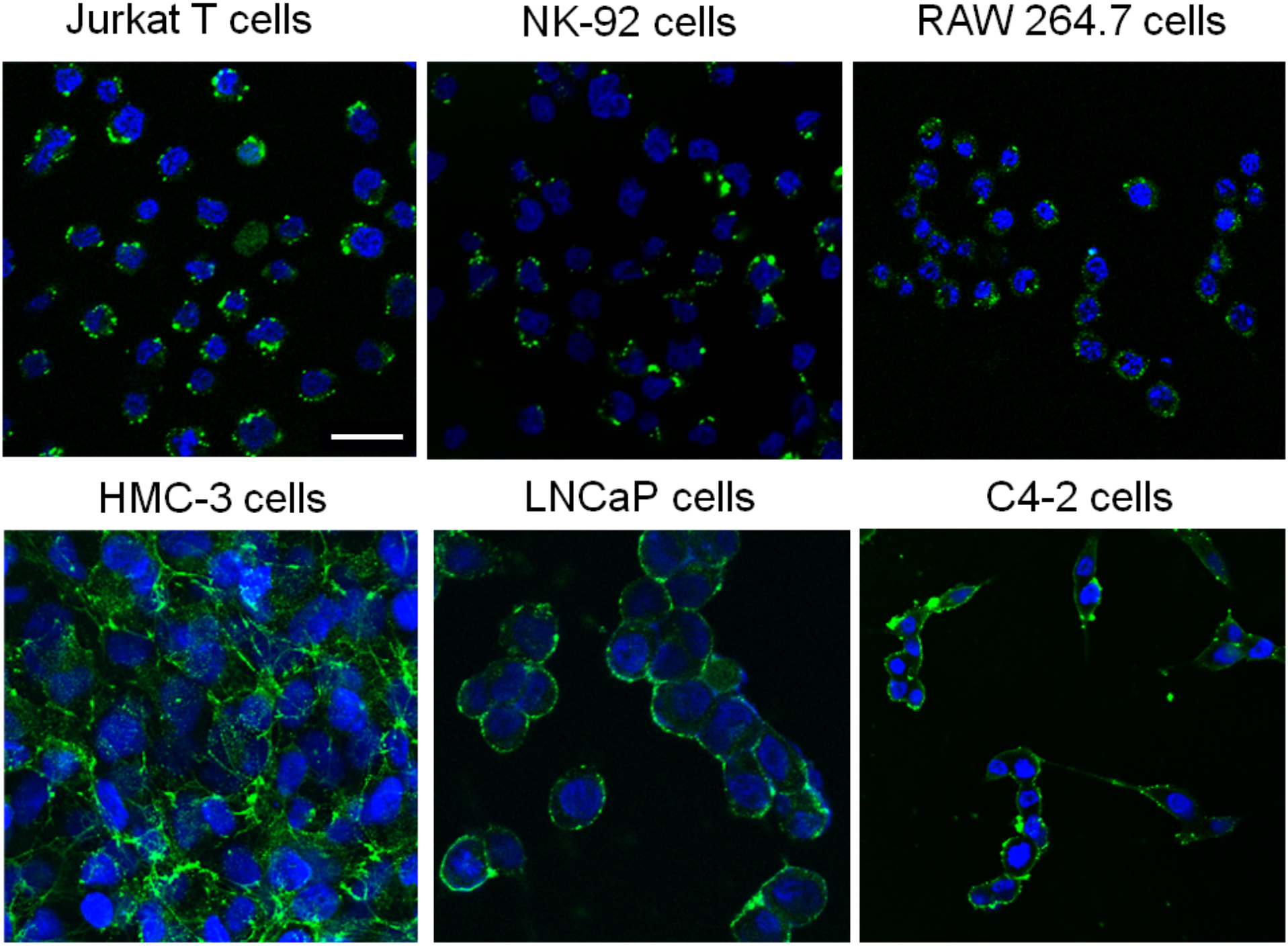
Live cell membrane imaging. Membrane imaging application of multiple live cells. Jurkat T cells, human natural killer NK-92 cells, mouse microphase RAW264.7 cells, human microglia HMC-3 cells, human prostate cancers cells (LNCaP and C4-2). Scale bar is 25 μm.

### Reversible and reusable reagent for live cell imaging

Finally, we were interested to study the reversibility and reusability of the dual-conjugated covalent cargo reagent for live cell imaging. We conjugated a magnetic bead to our dual-conjugated covalent cargo that was designed to form bonds with cell membrane such that the bonds are stimulated by pH. The magnetic beads were essential to isolate the cargo reagent, both from excess cargo that was not conjugated to cells at 0.1 mg/mL concentration during initial conjugation (unconjugated magnetic cargo), as well as, after cell-surface bond disruption and recovery of the cell conjugated cargo reagent by pH stimulation (pH recovered magnetic cargo), as shown in schematic representation in Figure 5a and Supplementary Figure S13. Covalent cargo conjugated Jurkat T cells were treated with PBS at pH 5.5 to disrupt surface conjugation, observed after 30 minutes of incubation, as evident from no green fluorescence on the cell surface. Next, the mixture of T cells and pH stimulated unconjugated magnetic cargo reagents were recovered using magnetic isolation from the supernatant. The pH recovered magnetic cargo solution was adjusted to pH 7.0 and reused in combination with the unconjugated magnetic cargo (Figure 5) to treat live Jurkat T cells. Surface conjugation of Jurkat T cells was observed for the combination of recovered cargo to test for reusability (Figure 5b, Supplementary Figure S14). To estimate the extent of reusability, we performed the surface conjugation-disruption-reconjugation several times. We observed that our cargo can be reused at least two times in reversible manner with the magnetically isolated reagent unlike a one-time use of existing reagents for live-cell imaging. Our dual-conjugated covalent cargo reagent can recover the cell culture with simple pH stimulated unconjugation and can be used multiple times for membrane imaging of live cells.

**Figure 5.**
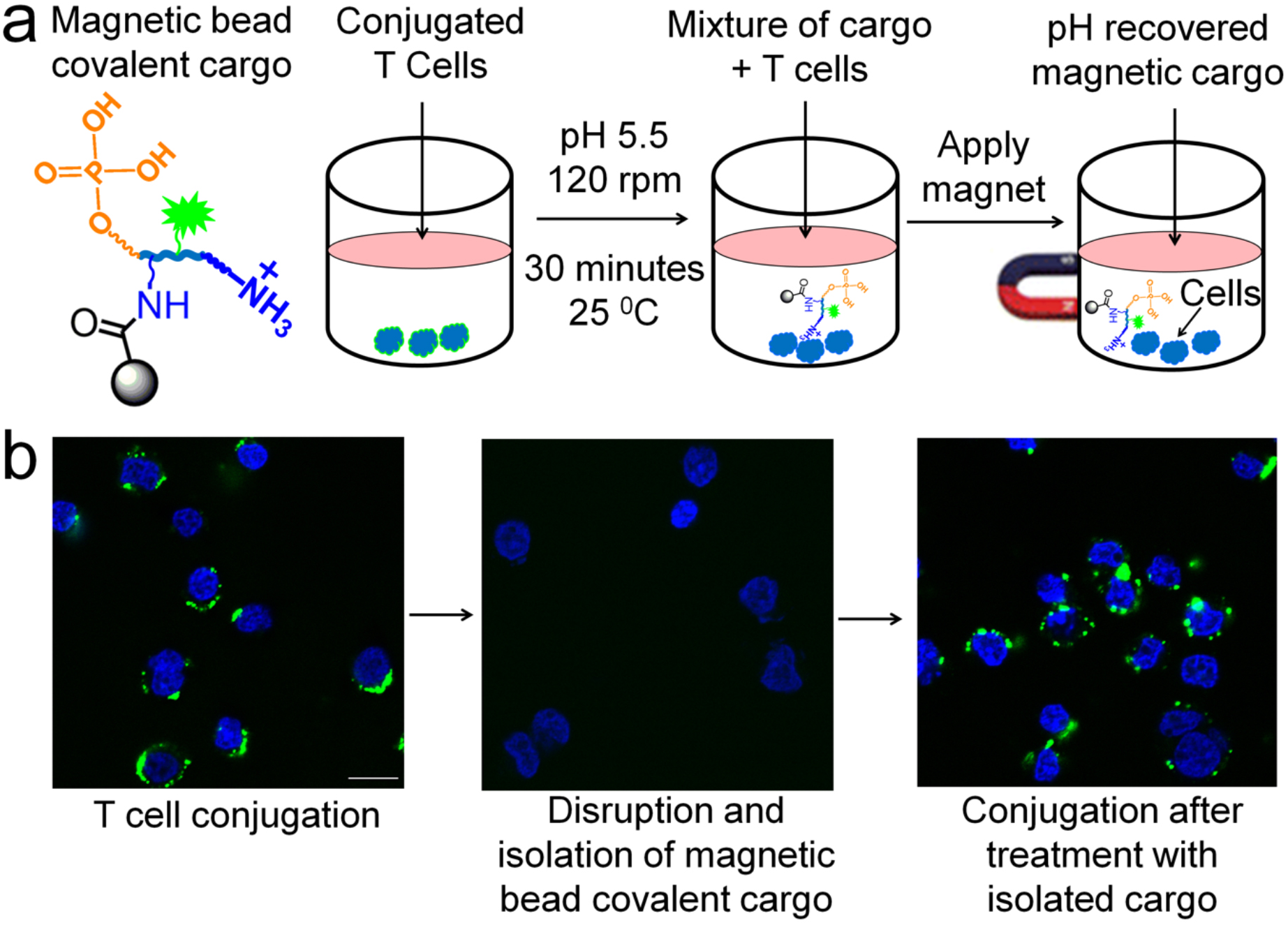
Reversibility and reusability of the cargo reagent for live cell imaging. (a) Structure of the magnetic bead linked covalent fluorophore cargo and schematic representation showing recovery of the magnetic cargo from surface conjugated T cells. (b) Confocal images of surface conjugated Jurkat T cells with covalent cargo reagent, after disruption and isolation of surface conjugation by pH stimulation, and then retreatment of isolated cargo reagent. Scale bar is 20 μm.

We have developed a cytocompatibility and stable method of chemical conjugation with live cell membranes. One side chain of our cargo molecule is a positively charged ammonium moiety, which provide electrostatic binding resulting in non-covalent conjugation with the cell surface; while the other side contains a phosphate group bearing ADP moiety that forms phospho-ester covalent conjugation with the cell surface phosphates. This dual conjugated modification strategy resulted in long-time stability with cell membrane and was non-toxic to live cells. While such an approach of chemical conjugation has many applications, we have shown one application of this method for live cell membrane imaging with long-time stability, recovery of the cell culture after imaging, and reusability of our reagent on live cells. Future work of this technology involves *in vivo* imaging and stability of our dual-conjugated cargo molecules that may be used to track therapeutic cells (e.g. CAR T-cells) *in vivo*, as well as, enhance the function of such primary cells for future clinical applications. Given the general conjugation chemistry exploited by our method, we expect several other applications of this dual-conjugation chemistry to functionalize membrane viruses, organisms (e.g. bacteria, yeast), etc. by conjugating with phospholipid moieties.

## Methods

### Materials

All the chemicals including NHS-fluorescein, adenosine di-phosphate (ADP), poly-d-lysine, phospholipid etc. were purchased from commercial suppliers and used without further purification. We purchased analytical grade solvents from commercial suppliers for synthesis. Fluorescence images of cells were captured using 40X and 60X objective using both Cytation 5 imaging reader and confocal laser microscope (Nikon AR1-MP) and 3D videos are generated as supporting information. Cell viability and proliferation assays was performed by using cell titer blue (CTB) reagent.

### Cell culture

We obtained Jurkat T cells from Dr. Majid Kazemian (Purdue University, USA) and NK-92 cells from Dr. Sandro Matosevic (Purdue University, USA). LNCaP, C4-2 and RAW 264.7 cell lines were provided by Professor Timothy Ratliff (Purdue University Center for Cancer Research, USA). HMC-3 cell line was given by Professor Jianming Li, Purdue University College of Veterinary Medicine. Cells were cultured following American Type Culture Collection (ATCC) protocol at 37 °C with 5% CO2 atmosphere in a humidified incubator. For normal growth, Jurkat T cells were cultured in RPMI-1640 media (Gibco) supplemented with 10% FBS (Atlanta Biologics), 20 mM HEPES and 1% penicillin/streptomycin (Invitrogen). NK-92 cells were cultured in RPMI-1640 supplemented with 100 IU/mL IL-2. All the cargo compounds were dissolved in PBS at high concentration (1 mg/mL) followed by filtrations using a 0.22 μm syringe filter and dilutions from this stock solution were prepared in culture medium.

### Synthesis of dual conjugated covalent cargo. Synthesis scheme for designed cargo is shown in Supplementary Figure S2

**Step 1.** Thionyl chloride (1.1 mmol) was added to a solution of benzotriazole (3 mmol) in dry DCM (10 mL) at room temperature and the reaction mixture was stirred for 10 min. Then iodoacetic acid (1.0 mmol) was added and the mixture was stirred for 12 h at room temperature. The white precipitate was filtered and concentrated under reduced pressure. After the evaporation of the solvent, the crude residue of benzotriazolide was isolated as intermediate and used in the next step reaction where it (0.06 mmol) was added to a HBr salt solution of D-lysine chain polymer (30 mg, 150 kD) in 5 mL of a MeOH in the presence of triethyl amine (100 μL). The reaction mixture was stirred at 4 °C for 24 h. After the evaporation of the solvent, crude residue of cargo intermediate I was isolated as precipitate which was purified by washing with acetonitrile and used in the next step.

**Step 2.** ADP (5.3 mg) was dissolved in 5 mL THF in a RB followed by addition of 4-(Dimethyl amino) pyridine (DMAP, 2.4 mg) to it. This mixture was stirred for 60 minutes at 4 °C followed by addition of cargo intermediate I (30 mg). The resulting reaction mixture was further stirred for 24 h at 4 °C. The reaction mixture was acidified with ice cold 1 (N) HCl to neutralize any remaining alkoxide of the ADP, as well as, form cationic ammonium chloride non-covalent probe in the cargo backbone. Finally, all the solvent was evaporated to get the crude product of covalent cargo which was purified by washing with acetonitrile.

### Synthesis of non-covalent phospholipid cargo

4.0 mg of HBr salt of D-lysine chain polymer (150 kD) was dissolved in 5 mL water. 10 μL triethyl amine was added and mixture was stirred for 5 minutes at room temperature (RT) to free all amine groups. Next, 2.0 mg phospholipid was dissolved in 2 mL methanol and added to it and the reaction mixture was stirred at RT. The reaction mixture was evaporated to get the crude residue and washed with acetonitrile to remove any un-reacted phospholipid and get the non-covalent phospholipid cargo.

### Synthesis of covalent phospholipid cargo

6.0 mg of covalent cargo was dissolved in 5 mL water followed by addition of 2.0 mg phospholipid and the reaction mixture was stirred at room temperature. The reaction mixture was evaporated at room temperature to get the crude residue, then washed with cold acetonitrile to remove any un-reacted phospholipid and get the title product.

### Synthesis of dual-conjugated covalent and non-covalent fluorophore cargo

Synthesis scheme for fluorophore conjugated cargo molecules is shown in Supplementary Figure S4. For dual-conjugated covalent cargo, 6.0 mg of covalent cargo prepared in the previous step (Supplementary Figure S2) was dissolved in 5 mL methanol by adding 100 μL of triethyl amine. This mixture was stirred for 30 minutes at 4 °C followed by addition of 1.0 mg NHS-Fluorescein. For non-covalent fluorophore cargo molecule, 8.0 mg HBr salt of D-lysine chain polymer was dissolved in 5 mL methanol by adding 100 μL triethyl amine. The resulting reaction mixture for each of the covalent and non-covalent cargo reactions (Supplementary Figure S4) was further stirred for 12 h at 4 °C. Un-reacted triethyl amine was neutralized by dropwise addition of ice cold 1 (N) HCl. The orange-yellow precipitate obtained from this reaction was separated and washed with cold methanol and dried at room temperature in vacuum to obtain the dual-conjugated fluorophore covalent cargo.

### Synthesis of butoxy carbonyl-protected non-covalent fluorophore cargo (BOC-cargo)

4.0 mg HBr salt of D-lysine chain polymer (150 kD) was dissolved in 5 mL methanol followed by adding 100 μL triethyl amine. This mixture was stirred for 30 minutes at 4 °C followed by addition of 2.0 mg NHS-fluorescein dye. The resulting reaction mixture was further stirred for 12 h at 4 °C. Next, excess BOC-anhydride (5.0 mg) was added to it and the reaction mixture was stirred for another 2 h. The precipitate obtained from this reaction was separated and washed with cold acetonitrile and dried at room temperature under vacuum to get the BOC-protected non-covalent fluorophore cargo.

### Synthesis of magnetic bead linked cargo

4.0 mg of each non-covalent and covalent fluorophore cargo was dissolved in 5 mL methanol by adding 100 μL triethyl amine. The reaction mixture was stirred for 30 minutes at 4 °C to make free all the amine groups. Next, 1.0 mg of NHS-magnetic beads was added to each of the reaction mixture and stirred for 12 h at 4 °C. Un-reacted triethyl amine was neutralized by dropwise addition of ice cold 1 (N) HCl. The brown-yellow precipitate obtained from this reaction was separated and washed with cold acetonitrile and dried at room temperature under vacuum to get the magnetic bead linked respective cargos.

### General procedure for cell-surface conjugation and live cell imaging

Jurkat T cells (100,000/well) were plated in a 12 well plate and treated with 0.1 mg/mL concentration of fluorescein, non-covalent and covalent fluorophore cargos in growth media. The cell and cargo mixture were shaken at 120 rpm using an orbital shaker for 30 minutes at room temperature. Next, cells were stained with 0.01 mg/mL concentration of Hoechst 33342 (for nucleus) in growth media and washed with sterile PBS and transferred in glass bottom dish. Cells were viewed under 60X oil object (optical zoom 3) in confocal laser microscope (Nikon AR1-MP).

### Stability of the conjugated cargo molecules on surface modified Jurkat T cells

Surface conjugated Jurkat T cells (100,000 cells/well) were grown in 12 well culture plate in growth media and images were captured after 1, 3 and 6 days of incubation using confocal laser microscope. Fluorescence intensity was measured using NIS-Elemental software.

### Legends for cell-conjugated 3D confocal movies

#### Movie 1: Surface conjugation of Jurkat T cell with covalent cargo

Jurkat T cells were treated with the covalent cargo (0.1 mg/mL) for 30 minutes in growth media at room temperature and stained with Hoechst 33342 for nucleus. Cells were then washed with PBS and confocal laser microscopic images were captured in z-stack using 60X oil object. Nucleus is shown in blue and cell surface is shown in green.

#### Movie 2: Surface conjugation of LNCaP prostate cancer cell with covalent cargo

LNCaP cells were seeded in glass bottom dish and grown in RPMI-1640 media for 70 % confluency. Cells were treated with the covalent cargo (0.1 mg/mL) for 30 minutes in growth media at room temperature followed by stained with Hoechst 33342 for nucleus. Finally, cells were washed with PBS and confocal images were captured using 60X oil object. Nucleus is shown in blue and cell surface is shown in green.

### Cargo displacement reactions with surface modified T cells with no fluorophore-tag cargo

Jurkat T cells were conjugated with the non-covalent and covalent fluorophore cargos for 30 minutes. Next, these surface conjugated Jurkat T cells (100,000 cells/well) were taken in 12 well culture plate and treated with no fluorophore tag non-covalent and the dual-conjugation covalent cargo for another 30 minutes in growth media at 120 rpm shaking at room temperature. Cells were treated with Hoechst 33342 for nuclear stain and washed with PBS. Finally, cells were transferred in glass bottom wells and confocal images were recorded to monitor the retention of surface conjugation.

### Viability assay of the surface modified cells

The cell viability experiment was performed using the CellTiter blue reagent. Surface conjugated Jurkat T cells (100,000 cells/well) were seeded in each well of 96-well plates using growth media and incubated in a humidified incubator at 37 °C and 5% CO_2_ atmosphere. At the end of the incubation, cell titer blue reagent (10 μl) was added directly to each well and the plates were incubated for additional 3 h at 37 °C to allow cells to convert resazurin to resorufin, and the fluorescent signal was measured at 590 nm after exciting at 560 nm using a multiplate ELISA reader (Bio-Tek Synergy HT plate reader, Bio-Tek, Winooski, VT). The percentage of live cells in a cargo-conjugated sample was calculated by considering the fluorescence intensity of the vehicle treated un-conjugated Jurkat T cell sample as 100%.

### Live cell membrane imaging

All the adherent cells (mouse microphase RAW264.7 cells, human microglia HMC-3 cells, human prostate cancers LNCaP and C4-2 cells) were plated and cultured on glass bottom dish following ATCC protocol. All these live cells were treated with 0.1 mg/mL of the covalent fluorophore cargo for 30 minutes in their respective growth media at RT at 120 rpm shaking. Cells were then stained with and washed with PBS. Finally, images were captured using 40X oil object in confocal laser microscope. For, both suspension Jurkat T and human natural killer NK-92 cells, general live cell imaging method was followed.

### Fixed cell surface conjugation and imaging

Jurkat T cells were mixed with 4 % fixing solution (4 % paraformaldehyde made in PBS) and immediately transferred in glass bottom dish. Cells were seeded as well as fixed by centrifugation at 1000 rpm at 10 °C for 5 minutes. Next, fixed cells were gently rinsed with PBS to remove any fixation agent and treated with the covalent fluorophore cargo (0.1 mg/mL) for 30 minutes in PBS at RT. Cells were stained with DAPI (4′,6-diamidino-2-phenylindole) and washed with PBS and again centrifuged to make sure their attachment on the glass bottom surface. Finally, confocal images were captured using 60X oil object.

### Reversibility and reusability of the covalent cargo reagent

Jurkat T cells were taken in 24 well plate and incubated with magnetic bead linked covalent fluorophore cargo reagent for 30 minutes at 120 rpm on an orbital shaker. The excess cargo and surface conjugated cell mixture was centrifuged at 500 rpm for 2 minutes to decant the ‘unconjugated cargo’ from the top of the well. Next, surface conjugated Jurkat T cells were taken in 24 well plate and incubated with PBS of pH 5.5 for 30 minutes at 120 rpm on an orbital shaker. The magnetic bead linked free cargo molecules were isolated by magnetic separation from its T cell mixture. The recovered cargo solution, thus obtained was adjusted to pH 7.0 by adding 1 (N) NaOH solution and re-sued with combination of unconjugated cargo to incubate live Jurkat T cells for another 30 minutes at 120 rpm. The precipitated Jurkat T cells obtained in each step were stained with Hoechst for nucleus and washed with PBS to remove any trace of cargo reagent. Confocal images were acquired using 60X oil object.

## Acknowledgements

Support from the Department of Chemistry at Purdue University, Ralph W. and Grace M. Showalter Research Trust, and the Purdue University Center for Cancer Research’s Jim and Diann Robbers Cancer Research Grant for New Investigators Award is gratefully acknowledged. We thank Professors Herman O. Sintim and David Thompson for critical reading and suggestions for the manuscript.

## Author contributions

GC designed research. JM performed researched and analyzed the data. JM and GC wrote the manuscript.

## Additional information

**Supplementary Information** containing NMR and FTIR spectra (Supplementary Figures S15-23) of the cargos and other supplementary Figures are available on the Publications website.

